# Transcriptional response modules characterise IL-1β and IL-6 activity in COVID-19

**DOI:** 10.1101/2020.07.22.202275

**Authors:** Lucy CK Bell, Cem Meydan, Jacob Kim, Jonathan Foox, Daniel Butler, Christopher E. Mason, Sagi D. Shapira, Mahdad Noursadeghi, Gabriele Pollara

**Author notes:** **Corresponding author** Dr Gabriele Pollara, Division of Infection & Immunity, Cruciform Building, Gower Street, London, WC1E 6BT, United Kingdom,. Twitter: @gpollara.

## Abstract

Dysregulated IL-1β and IL-6 responses have been implicated in the pathogenesis of severe Coronavirus Disease 2019 (COVID-19). Innovative approaches for evaluating the biological activity of these cytokines *in vivo* are urgently needed to complement clinical trials of therapeutic targeting of IL-1β and IL-6 in COVID-19. We show that the expression of IL-1β or IL-6 inducible transcriptional signatures (modules) reflects the bioactivity of these cytokines in immunopathology modelled by juvenile idiopathic arthritis (JIA) and rheumatoid arthritis. In COVID-19, elevated expression of IL-1β and IL-6 response modules, but not the cytokine transcripts themselves, is a feature of infection in the nasopharynx and blood, but is not associated with severity of COVID-19 disease, length of stay or mortality. We propose that IL-1β and IL-6 transcriptional response modules provide a dynamic readout of functional cytokine activity *in vivo*, aiding quantification of the biological effects of immunomodulatory therapies in COVID-19.

## Introduction

Severe Coronavirus Disease 2019 (COVID-19) typically occurs over a week from symptom onset, when viral titres have diminished, suggesting a dysregulated host inflammatory response may be driving the pathogenesis of severe disease (Bullard et al., 2020; Huang et al., 2020; McGonagle et al., 2020). Elevated IL-1β and IL-6 responses have each been associated with disease severity (Huang et al., 2020; Liao et al., 2020; Qin et al., 2020; Ravindra et al., 2020; Zhang et al., 2020; Zhou et al., 2020b). In addition, the hyperinflammatory state in COVID-19 is reported to resemble some aspects of haemophagocytic lymphohistiocytosis (HLH), a condition that may benefit from therapeutic IL-1β blockade (Mehta et al., 2020). These observations have generated hypotheses that IL-1β and/or IL-6 may be key drivers of pathology in severe COVID-19, and led to clinical trials of IL-1β and IL-6 antagonists in this context (Maes et al., 2020). Randomised studies to date investigating the role of tocilizumab, a humanised monoclonal antibody against the IL-6 receptor, have shown no clinical benefit, but immunophenotyping beyond the measurement of single cytokines, before or after drug administration, was not recorded or correlated with clinical responses at the individual patient level (Hermine et al., 2020; Salvarani et al., 2020; Stone et al., 2020).

The measurement of individual cytokines at the protein or RNA level may not reflect their biological activity accurately within multivariate immune systems that incorporate redundancy and feedback loops. To address this limitation, we have previously derived and validated gene expression signatures, or modules, representing the transcriptional response to cytokine stimulation, using them to measure functional cytokine activity within genome-wide transcriptomic data from clinical samples (Bell et al., 2016; Byng-Maddick et al., 2017; Dheda et al., 2019; Pollara et al., 2017). However, transcriptional modules to quantify IL-1β or IL-6 response have not been used in COVID-19 to quantify the bioactivity of these cytokine pathways *in vivo*. In the present study, we have sought to address this gap, describing the derivation and validation of IL-1β and IL-6 inducible transcriptional modules, and testing the hypothesis that these modules can be used in the molecular assessment of the pathophysiology and the response to therapeutic cytokine blockade of inflammatory conditions, including COVID-19.

## Results

### Identification and validation of IL-1β and IL-6 transcriptional modules

We first sought to derive transcriptional modules that identified and discriminated between the response to IL-1β and IL-6 stimulation. We have previously derived an IL-1β response module from cytokine stimulated fibroblasts (table S2) (Pollara et al., 2019). As in our prior studies (Bell et al., 2016; Pollara et al., 2017, 2019), we used the geometric mean of the constituent genes in a module as a summary statistic to describe the relative expression of the module. We demonstrate that in both monocyte-derived macrophages (MDM) and peripheral blood mononuclear cells (PBMC) (Boisson et al., 2012; Jura et al., 2008), IL-1β stimulation induced greater expression of the IL-1β response module than either IL-6 or TNF_α_ stimulation, where there was no increased expression above unstimulated cells (Fig 1A + B). To identify an IL-6 response module which was able to discriminate from the effects of IL-1β, we identified one study that had stimulated human MDM with either IL-1β (15 ng/ml) or IL-6 (25 ng/ml) for 4 hours (Jura et al., 2008). Hierarchical clustering identified genes induced by IL-6 but not IL-1β, and we termed this the IL-6 response module (table S2). Internal validation of this module confirmed increased expression in IL-6 stimulated MDM (fig 1A). Testing the IL-6 module in other datasets demonstrated elevated expression following IL-6, but not TNF_α_, stimulation of human kidney epithelial and macrophage cell lines (Das et al., 2020; O’Brown et al., 2015) (figs 1C+D), whereas no elevated expression of the IL-6 module was observed following IL-1β or TNF_α_ stimulation of MDM or PBMC (figs 1A+B). These findings demonstrated that the IL-1β and IL-6 response modules could detect the effects of their cognate cytokines, and discriminate these from each other and from an alternative inflammatory cytokine stimulus, TNF_α_.

**Figure 1.**
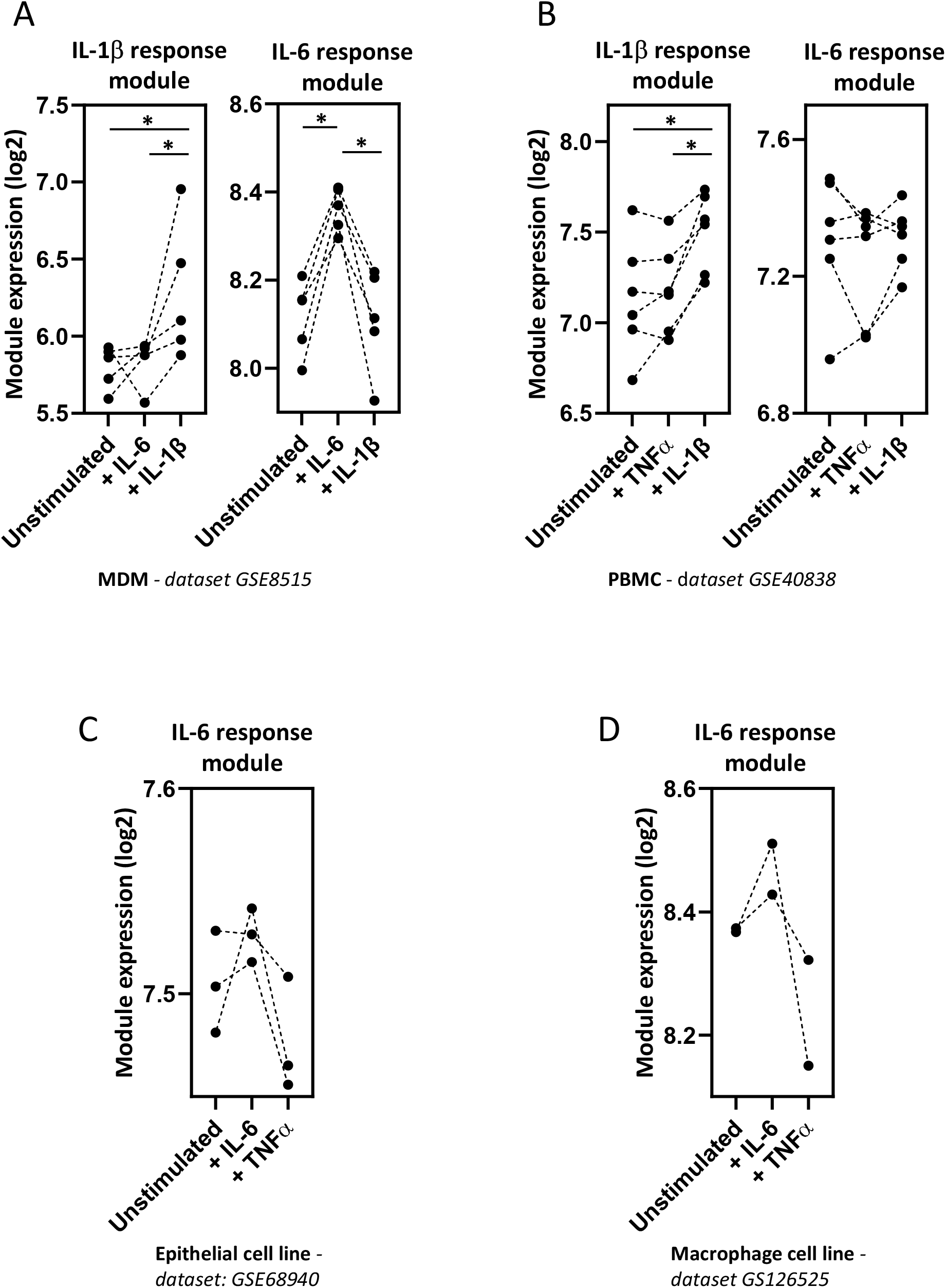
Validation of cytokine response modules. Geometric mean module expression in A) MDM stimulated *in vitro* with either IL-1β (15 ng/ml) or IL-6 (25 ng/ml) for 4 hours (Jura et al., 2008), B) PBMC stimulated with TNF_α_(20 ng/ml) or IL-1β (10 ng/ml) for 6 hours (Boisson et al., 2012), C) human renal proximal tubular epithelial (HK-2) cells stimulated with IL-6 (200 ng/ml) or TNF_α_(100 ng/ml) for 1.5 hours (O’Brown et al., 2015) and D) human macrophage cell lines (THP-1) stimulated with IL-6 (50 ng/ml) or TNF_α_(10 ng/ml) for 2 hours (Das et al., 2020). Transcriptomic datasets are designated adjacent to figure panels. * = p < 0.05 by Mann-Whitney test.

### IL-1β and IL-6 module expression in chronic inflammation

To determine whether IL-1β and IL-6 response modules were able to detect elevated cytokine bioactivity *in vivo*, we assessed the blood transcriptome of juvenile idiopathic arthritis (JIA) and rheumatoid arthritis (RA) patients. These are conditions in which elevated IL-1β and IL-6 activity are considered to play a key role in disease pathogenesis, evidenced by clinical improvement following therapeutic antagonism of these cytokines (De Benedetti et al., 2012; Fleischmann, 2017; Nikfar et al., 2018; Ruperto et al., 2012). The blood transcriptome of untreated JIA patients displayed elevated IL-1β and IL-6 bioactivity (fig 2A) (Brachat et al., 2017), but this was not consistently evident in several RA blood transcriptome datasets (fig S1) (Lee et al., 2020; Macías-Segura et al., 2018; Tasaki et al., 2018). Discrepancies between molecular changes in blood and tissues have been previously described in RA (Lee et al., 2020), and therefore we tested the hypothesis that in contrast to blood, elevated IL-1β and IL-6 bioactivity was a feature of the synovium in RA. Consistent with this hypothesis, a separate transcriptomic dataset of synovial membrane biopsies from patients with RA (Broeren et al., 2016) showed elevated levels of both IL-1β and IL-6 response module expression compared to non-RA synovium (fig 2B).

**Figure 2.**
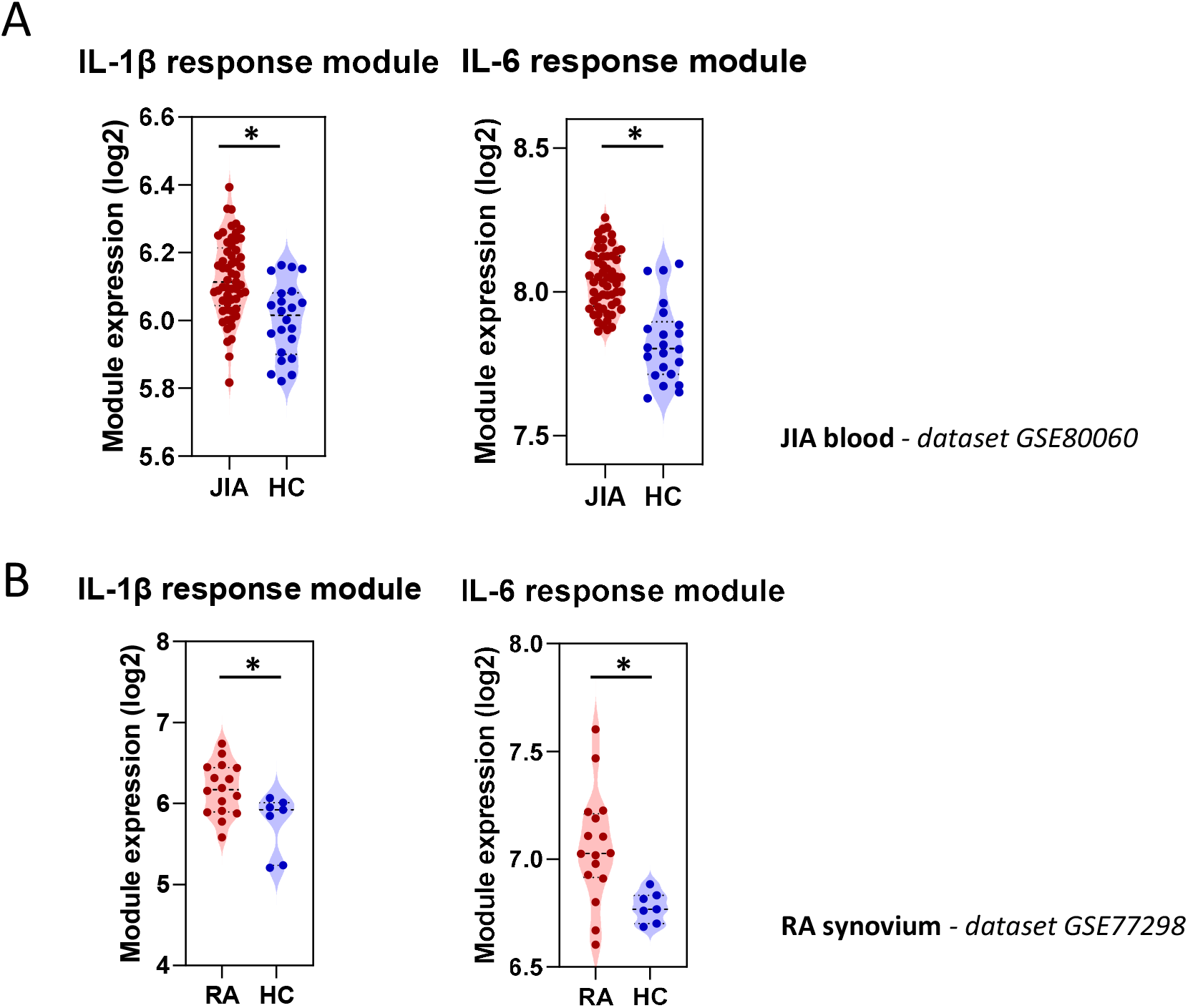
Cytokine response module expression in chronic inflammatory conditions. Geometric mean expression of IL-1β and IL-6 cytokine response modules in A) blood of patients with JIA compared to healthy controls (Brachat et al., 2017), and B) in the synovium of RA patients compared to that of healthy controls (Broeren et al., 2016). Transcriptomic datasets are designated adjacent to figure panels. * = p < 0.05 by Mann-Whitney test.

We used the elevated cytokine activity in the blood of JIA patients to test the hypothesis that therapeutic cytokine modulation would result in changes in cytokine bioactivity as determined by module expression. We made use of the blood transcriptome of JIA patients 3 days following administration of canakinumab, a human monoclonal antibody to IL-1β (Brachat et al., 2017). Patients who had a therapeutic response to canakinumab showed elevated IL-1β module expression which reduced 3 days after canakinumab administration (fig 3A). In contrast, in those who had no treatment response, IL-1β module expression was lower at baseline and was unaffected by canakinumab (fig 3A). Unlike the differences seen in the IL-1β module between responders and non-responders, there were no differences between these groups in IL-6 module expression at baseline (fig 3B). This indicated that these two cytokine response modules quantified two distinct biological processes. Interestingly, expression of the IL-6 module was also diminished after canakinumab treatment in patients who responded to treatment, suggesting that IL-6 activity may be downstream of IL-1β in this context. Of note in these populations, the expression of the *IL1B* gene correlated with that of the IL-1β response module, but the same was not evident between IL-6 module and *IL6* gene expression (fig 3C), illustrating an example in which cytokine gene expression itself may not necessarily reflect the functional activity of that cytokine.

**Figure 3.**
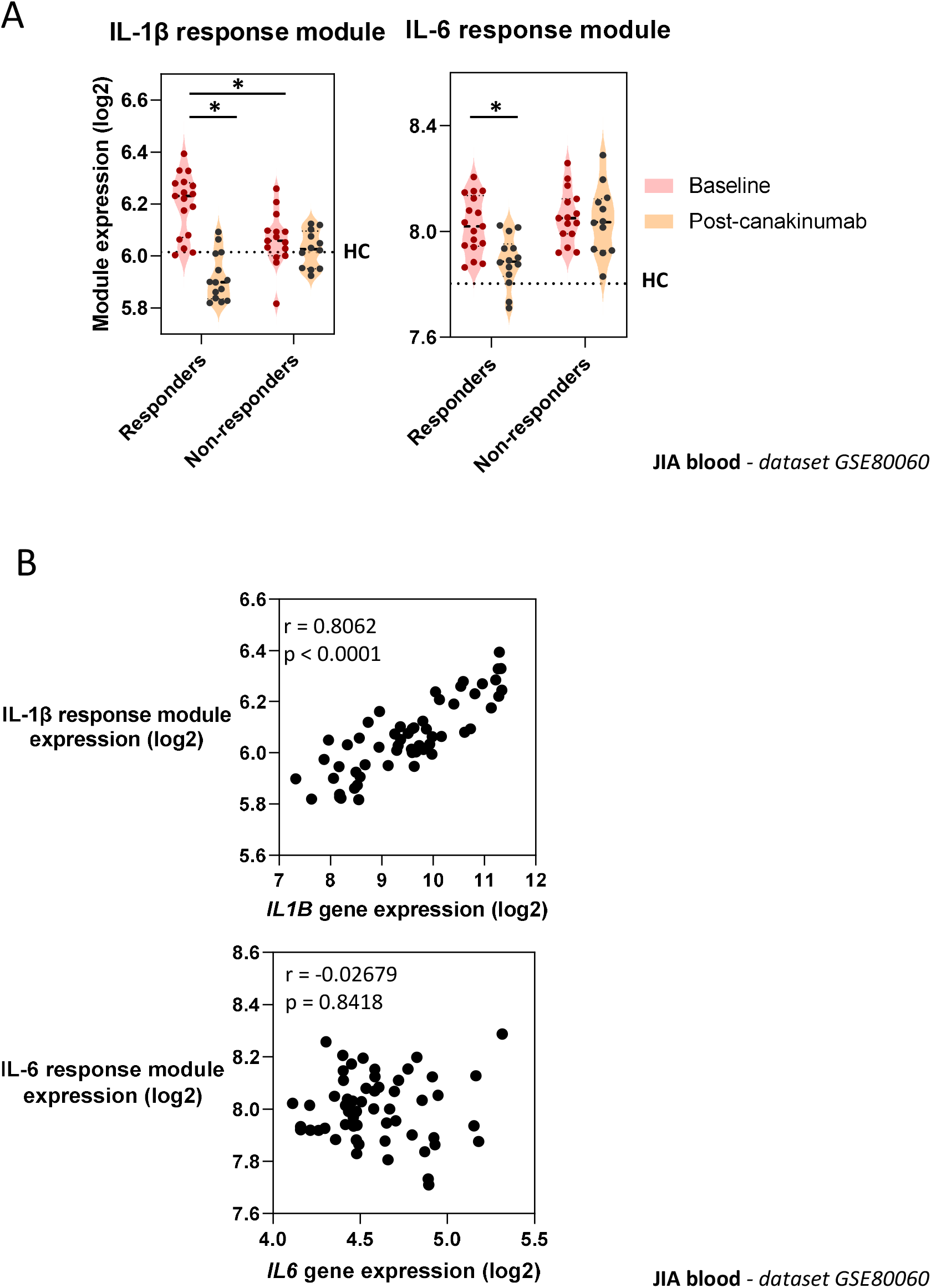
Effect of canakinumab on expression of cytokine response modules and genes. A) Geometric mean expression of IL-1β and IL-6 cytokine response modules in JIA patients before and 3 days after administration of canakinumab (Brachat et al., 2017). Patients were subdivided into good responders (90-100% improvement) and non-responders (0-30% improvement). Dotted lines indicate median module or gene expression in healthy controls (HC) population in same dataset. * = p < 0.05 by Mann-Whitney test. B) Relationship between expression of cytokine response modules and cytokine genes. Statistical assessment of correlation made by Spearman Rank correlation. r = correlation coefficient. Transcriptomic dataset designated adjacent to figure panels.

### IL-1β and IL-6 bioactivity in COVID-19

We tested the hypothesis that elevated IL-1β and IL-6 bioactivity is a feature of COVID-19 disease. We initially explored the induction of IL-1β and IL-6 activity at the site of COVID-19 disease, by profiling transcriptional responses in nasopharyngeal swabs from 495 control and 155 SARS-CoV-2 infected individuals (Butler et al., 2020; Ramlall et al., 2020). Gene set enrichment analysis (GSEA) was used as an alternate method of module enrichment scoring (Subramanian et al., 2005), in line with previous analyses of this data set (Ramlall et al., 2020). While the IL-1β response module was modestly induced by SARS-CoV-2 infection, the IL-6 response module was significantly enriched in transcriptional programs induced by this viral infection (fig 4). Moreover, we found that SARS-CoV-2 viral loads were positively associated with cytokine activity, with enrichment of IL-1β and IL-6 responses observed in individuals with the upper tertile of measured viral loads, while patients with the lowest tertile viral titres did not show induction of responses to either cytokine (fig 4). The greatest IL-6 responses were in fact observed in individuals with intermediate viral titres, in whom significant induction of IL-1β activity was not seen (fig 4). Together, these findings suggest that both IL-1β and IL-6 activity are a feature of the host response at the site of SARS-CoV-2 infection, and are likely to be driven by increasing viral replication *in vivo*.

**Figure 4.**
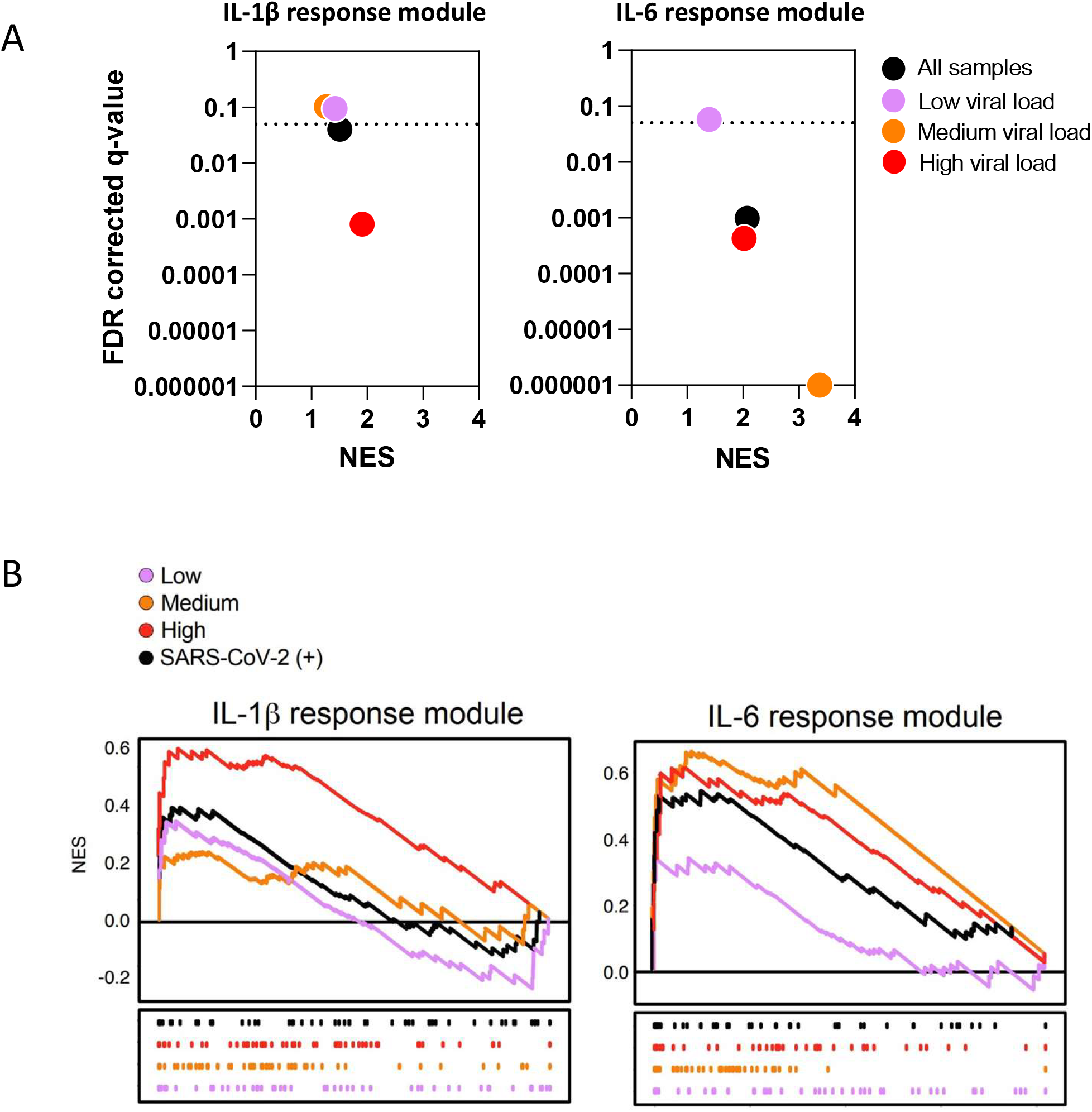
Cytokine response modules at the site of disease in COVID-19. A) Gene set enrichment analysis (GSEA) of the IL-1β and IL-6 modules was applied to nasopharyngeal swabs from SARS-CoV-2 infected and uninfected individuals. Patients were stratified into low (pink), medium (orange) and high (red) viral loads as previously described (Ramlall et al., 2020). GSEA was used to determine the level of engagement for the respective modules in the context of SARS-CoV-2 infection (Subramanian et al., 2005), in line with previously published analysis of this data set (Ramlall et al., 2020). Normalised enrichment scores (NES) are shown on the x axes and measurement of statistical significance (false detection rate q-value) is shown on the y axes. The threshold for significance (q=0.05) is shown by the dotted lines; data points below the dotted lines are significantly enriched for the relevant module in each group of SARS-CoV-2 positive patients, in comparison to the control group. B) Leading edge enrichment plots from GSEA of the cytokine modules for each comparison.

As clinical deterioration in COVID-19 occurs after peak viral replication in the airways has subsided, we tested the hypothesis that IL-1β and IL-6 activity was also related to disease severity. We initially explored IL-1β and IL-6 activity in the blood of 3 patients with mild-moderate COVID-19 disease who were admitted to hospital and recovered (Ong et al., 2020). This dataset was generated using the Nanostring system and consisted of 579 mRNA targets, which included only 7/57 (12.2%) and 7/41 (17.1%) constituent genes of the IL-1β and IL-6 response modules respectively (table S2). We demonstrated that IL-1β and IL-6 submodules, generated from these shorter lists of constituent genes, were still able to recapitulate all the findings from fig 3 (fig S2). The expression of these submodules in the blood transcriptome of this small number of COVID-19 patients revealed variation in IL-1β and IL-6 bioactivity over the period of hospitalisation, with higher expression seen earlier during hospital admission and a reduction as patients recovered (Fig 5A). This time-associated relationship with clinical recovery was not seen for the expression of the *IL1A*, *IL1B* and *IL6* genes (fig 5A). We extended these analyses by assessing the transcriptome of blood samples collected at the time of hospital admission from 32 COVID-19 patients presenting with varying levels of disease severity (Hadjadj et al., 2020). These data, also collected using the Nanostring system, revealed expression of the IL-1β and IL-6 cytokine submodules was clearly elevated in COVID-19 compared to healthy controls (fig 5B). However, strikingly, there was only minimal variability in IL-1β and no variability in IL-6 submodule expression between the different levels of COVID-19 disease severity (fig 5B).

**Figure 5.**
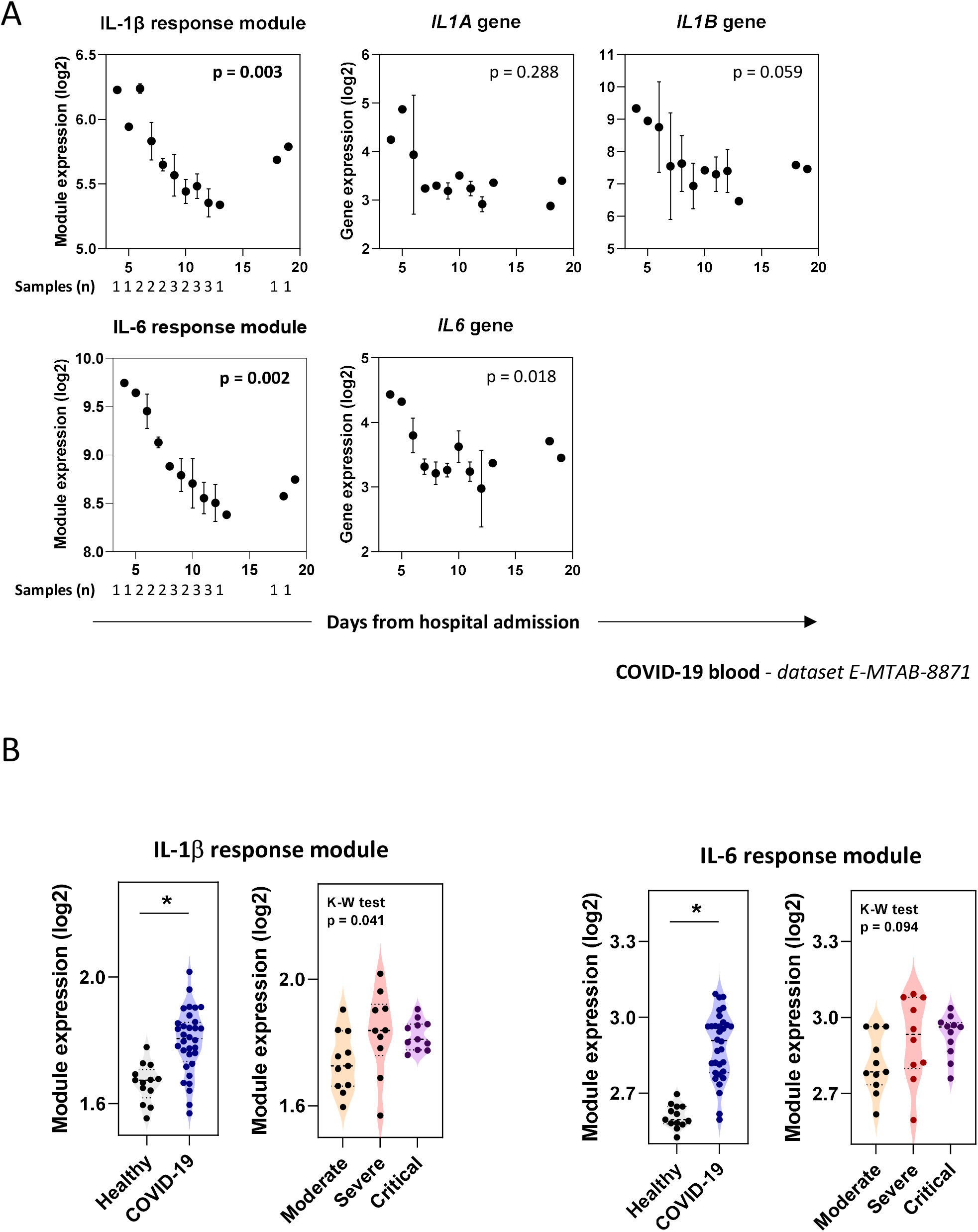
Cytokine response module and gene expression in COVID-19 blood samples. A) Geometric mean expression of IL-1β and IL-6 response module and *IL1A*, *IL1B* and *IL6* gene expression in patients admitted with COVID-19 (Ong et al., 2020). Number of patient samples at each timepoint designated on first plot of each row, but applicable for all panels. Where more than one sample was available at any time point, the mean expression +/− SEM is plotted. Kruskal-Wallis test was performed on binned time points 4-6, 7-9, 10-12 and 12+ days following hospitalisation, corresponding to 4, 7, 8 and 3 samples in each of these categories. The p values shown represent Kruskal-Wallis tests with time since hospital admission as the independent variable, where a threshold of 0.01 (corrected for multiple testing by the Bonferroni method) is required for a single test to be classed as significant (significant p-values indicated in bold text). B) Geometric mean expression of IL-1β and IL-6 response modules in whole blood transcriptomic profiles from patients admitted with moderate (n=11) severe (n=10) or critical (n=11) COVID-19, in comparison to healthy controls (n=13) (Hadjadj et al., 2020). In this study, samples were collected from patients at the time of admission to hospital, a median of 10 days (IQR 9 – 11 days) from symptom onset. A Mann-Whitney test was used to assess differences in module expression between all COVID-19 patients and healthy controls (* = p < 0.05), and a Kruskal-Wallis test was used to determine variability in module expression between the grades of COVID-19 disease severity.

Finally, we tested the hypothesis that elevated IL-1β and IL-6 transcriptional activity in blood could predict clinical outcome in COVID-19. We assessed the transcriptome of blood leucocytes from 101 COVID-19 and 24 non-COVID-19 patients admitted to hospital (Overmyer et al., 2020). As seen in the whole blood transcriptome analysis (fig 5), leucocytes from COVID-19 patients also demonstrated elevated IL-1β and IL-6 module activity compared to controls (fig 6A), and once again this distinction was not seen in IL1A, IL1B and IL6 gene expression (fig S3). Clinical outcome in this cohort was determined from the number of hospital free days at day 45 (HFD-45) following hospital admission, whereby zero days indicated continued admission or death (Overmyer et al., 2020). Prognostication models have identified decreased lymphocyte counts as predictors of clinical deterioration (Gupta et al., 2020). Focusing on COVID-19 patients not requiring ICU admission, we reproduced this observation, demonstrating a positive correlation between HFD-45 and the expression of a transcriptional module that reflects T cell frequency *in vivo* (Pollara et al., 2017) (fig 6B). In contrast, neither IL-1β nor IL-6 response module expression at the time of study recruitment was associated with HFD-45, indicating that, in this dataset, transcriptional activity of these cytokines was not predictive of clinical outcome from COVID-19 infection (fig 6B).

**Figure 6.**
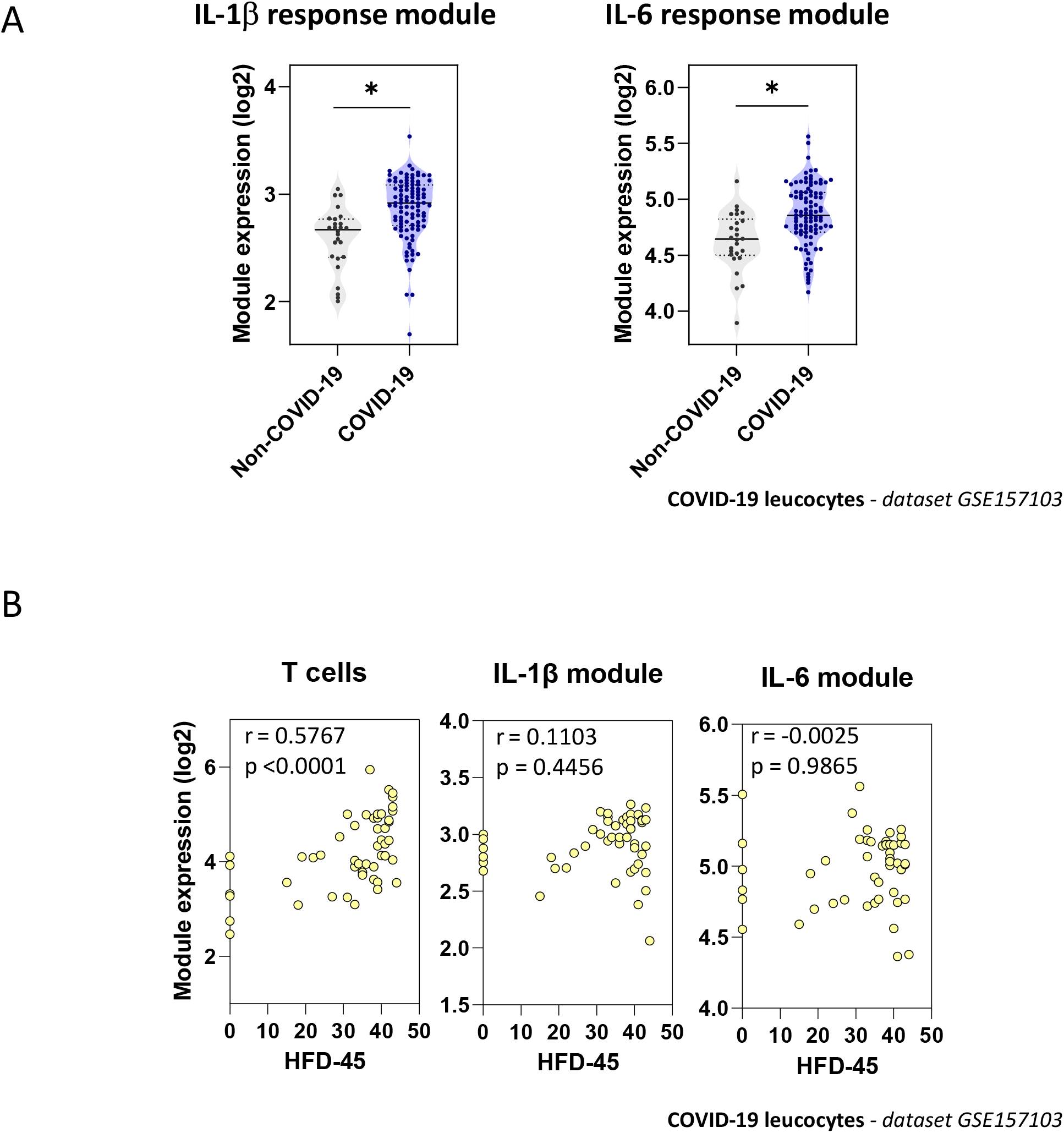
Relationship between cytokine response module expression at admission in COVID-19 and clinical outcome. A) Geometric mean expression of IL-1β and IL-6 response modules in transcriptomic profiles of blood leucocytes collected from 101 COVID-19 and 24 non-COVID-19 patients. In this study, samples were collected from patients at a median of 3.37 days from admission to hospital (Overmyer et al., 2020). B) In patients from this cohort who were not admitted to ITU, the relationship between expression of cytokine response modules, or a previously validated T-cell module (Pollara et al., 2017), and the number of hospital free days at day 45 (HFD-45) following hospital admission (whereby zero days indicated continued admission or death) is shown. Statistical assessment of correlation made by Spearman Rank correlation. r = correlation coefficient.

## Discussion

The protracted clinical course, inverse relationship between viral load and symptom progression, and the association between inflammation and worse clinical outcomes support a hypothesis whereby severe COVID-19 disease is predominantly driven by an exaggerated inflammatory response (Bullard et al., 2020; Huang et al., 2020). Both IL-1β and IL-6 may play a role in this process (Huang et al., 2020; Liao et al., 2020; Qin et al., 2020; Ravindra et al., 2020; Zhang et al., 2020; Zhou et al., 2020a), and cytokine modulating therapies are now being tested in COVID-19 clinical trials. In this study we utilised transcriptional modules derived from cytokine stimulated cells to demonstrate that their expression, but not that of their cognate cytokine genes, provided a quantitative readout for cytokine bioactivity *in vivo*, both in the context of COVID-19 and chronic inflammatory conditions.

We show that in COVID-19, IL-1β and IL-6 cytokine activity is detectable at a site of disease, the nasopharynx, where greater IL-6 bioactivity in particular is associated with higher levels of SARS-CoV-2 detected. This finding indicates that the presence of viral antigen is associated with IL-6 mediated inflammation, although we cannot ascertain from these experiments whether IL-6 inflammation persists in tissues in the later stages of severe COVID-19 when viral titres diminish (Bullard et al., 2020). The elevated cytokine responses seen in nasopharyngeal tissues were also detectable in the transcriptome of whole blood and isolated leucocytes from COVID-19 patients compared to the control populations available, although this analysis merits being extended to include a wider array of conditions associated with hyperinflammation (Leisman et al., 2020). Although a reduction in cytokine activity tracked clinical recovery from illness, IL-1β and IL-6 activity at the time of hospital attendance was not predictive for clinical outcome, and, in contrast to the association seen with circulating levels of IL-6 protein (Thwaites et al., 2020), we observed no clear gradient of IL-1β or IL-6 response module expression with disease severity. Our findings may help explain the recent results from randomised studies whereby neutralisation of IL-6 activity by tocilizumab did not show a benefit in mortality or clinical recovery in patients with severe COVID-19 (Hermine et al., 2020; Salvarani et al., 2020; Stone et al., 2020). However, these studies did not record IL-6 activity before or after tocilizumab administration, precluding associations between cytokine activity, neutralisation efficiency and clinical outcomes. We propose that future randomised trials will need to incorporate assessments of cytokine activity in study protocols to permit mechanistic correlations between immunomodulatory interventions and disease outcomes, promoting a stratified medicine approach to host-directed therapies in COVID-19.

A consistent observation in our work was that transcriptional modules identified differences between patient groups that would not otherwise have been detected by assessment of cognate gene transcripts. An interpretation of these findings is that the downstream response to cytokine stimulation is more persistent than the expression of the cytokine gene mRNA, the stability of which is subject to trans-regulatory factors and feedback loops (Iwasaki et al., 2011; Seko et al., 2006). Moreover, transcriptional modules are intrinsically composed of genes with co-correlated expression, minimising technical confounding of single gene measurements, demonstrated by the strongly concordant expression between the full and Nanostring subset IL-1β and IL-6 response modules. These factors may explain the discordance recorded between IL-6 gene expression and protein secretion in COVID-19 (Hadjadj et al., 2020). Moreover, cytokine levels after modulation *in vivo* do not necessarily reflect bioactivity, exemplified by the rise in IL-6 in blood following administration of tocilizumab (Nishimoto et al., 2008). We propose that cytokine response modules overcome both issues by integrating the culmination of cytokine signalling events, and may be used as an *in vivo* biomonitor of cytokine activity (Hedrick et al., 2020).

Our study has limitations. Despite drawing on four independent COVID-19 datasets, the sample sizes assessed in our study were still modest, especially for longitudinal samples, but this was limited by the data available. Assessments of the transcriptome from leucocytes and whole blood in COVID-19 may not be interchangeable and will need cross-validating, although both datasets demonstrated no association between IL-1β or IL-6 activity and severity of disease. Determining the sensitivity and specificity of the IL-1β and IL-6 response modules for their respective cognate cytokines was limited by the available datasets and the range of cytokine stimulation conditions performed in those experiments. Comparing the expression of these modules across a wider range of biologically paired cytokine stimulations will allow refinement of their accuracy. As the modules were generated from *in vitro* experiments, we sought to determine their applicability *in vivo*, assessing neutralisation of cytokine activity following immunomodulation with biologic agents *in vivo*. IL-1β activity in blood and in tissues was diminished days after canakinumab (fig 3) and anakinra (Pollara et al., 2019) administration respectively, but no equivalent datasets were available to assess the IL-6 response module in the same manner. Biobanked samples from ongoing tocilizumab clinical trials in COVID-19 and other diseases may provide an opportunity to validate IL-6 module performance in this way.

In conclusion, our data demonstrate elevated activity of the inflammatory cytokines IL-1β and IL-6 in COVID-19 in blood and tissues, and demonstrate the utility of cytokine transcriptional response modules in providing a dynamic readout of the activity of these pathways *in vivo*. We propose that use of these modules may enhance efforts to investigate the pathology of COVID-19, support development of methods to stratify patients’ risk of clinical progression, and aid quantification of the biological effects of host-directed immunomodulatory therapeutics in COVID-19.

## STAR methods

### Datasets

All datasets used are provided in table S1. Data matrices were obtained from processed data series downloaded from the NCBI Gene Expression Omnibus (GEO) (https://www.ncbi.nlm.nih.gov/geo/) or Array Express repository (https://www.ebi.ac.uk/arrayexpress/). Probe identifiers were converted to gene symbols using platform annotations provided with each dataset. In circumstances where downloaded datasets were not log_2_ transformed, this was performed on the entire processed data matrix. Duplicate genes were removed after the first one identified using Microsoft Excel duplicate remover function.

### IL-1β and IL-6 module derivation

We previously derived an IL-1β transcriptional module from the transcriptome of fibroblasts stimulated with IL-1β or TNF_α_ (Pollara et al., 2019). We derived a novel IL-6 transcriptional response module from a publicly available dataset (table S1) reporting experiments of human monocyte-derived macrophages (MDM) stimulated with IL-1β (15 ng/ml) or IL-6 (25 ng/ml) for 4 hours (Jura et al., 2008). In this study the transcriptional programme of cytokine-stimulated MDM was assessed by microarrays and hierarchical clustering was performed using Euclidean distance and average linkage method. This approach identified several unique clusters of genes differentially expressed following stimulation with each cytokine. Genes in clusters D & E showed elevated expression following stimulation by IL-6, but not by IL-1β. We combined the list of genes within these two clusters, removed duplicate or non-annotated genes, and termed this the IL-6 response module.

We applied the above IL-1β and IL-6 response modules to two studies where transcriptional profiling was performed using the Nanostring nCounter Human Immunology_v2 panel, which assesses the expression of a subset of the whole genome (579 genes) (Hadjadj et al., 2020; Ong et al., 2020). Consequently, only a subset of the modules’ constituent genes was present in these datasets (table S2). To verify the validity of applying our method to these datasets, we generated new cytokine response submodules using only genes from this subset, and showed them to provide the same discrimination of IL-1β and IL-6 responses as the parent modules (fig S2).

### Module expression assessment

The expression of transcriptional modules was derived by calculating the geometric mean expression of all constituent genes, as previously described (Pollara et al., 2017). The scripts used allowed the absence of a constituent gene in the analysed dataset, a scenario that did not affect geometric mean calculation. Gene set enrichment analysis was also used for modular expression assessment in nasopharyngeal samples, as previously described (Ramlall et al., 2020; Subramanian et al., 2005).

### Statistical analysis

All module score calculations were calculated in R v3.6.1 and RStudio v1.2.1335, using scripts generated and deposited in our previous publication (https://github.com/MJMurray1/MDIScoring) (Pollara et al., 2017). Mann-Whitney tests, Spearman rank correlations and Kruskal-Wallis tests were calculated in GraphPad Prism v8.4. Kruskal-Wallis testing was chosen to determine the presence of variability in the expression of cytokine response modules or cytokine gene over time since hospital admission (fig 5A) or between different categories of COVID-19 disease severity (fig 5B). This non-parametric test was chosen as we could not assume the expression of these variables was Normally distributed. In fig 5A patient samples were aligned according to days from hospital admission, and then binned into day interval categories (4-6, 7-9, 10-12 and 12+ days following admission), yielding 4, 7, 8 and 3 samples in each group. Kruskal-Wallis testing was performed on these binned categories to identify variation in the expression of modules or genes between these categories, with the Bonferroni method used for multiple testing correction.

### Role of funders

The funding sources played no role in conceiving the study, performing data analyses, preparing the manuscript or deciding to submit it for publication.

### Ethics statement

The manuscript makes use of publicly available datasets, the use of which required no further ethical approval.

## Author contributions

LCKB, MN and GP conceived the study. LCKB, CM, JK, JF, DB, CEM, SDS and GP performed the analyses. LCKB, SDS, MN and GP critically appraised the results, drafted the manuscript, and agreed on the data presented and the conclusions reached in the final version. All authors reviewed and approved the manuscript.

## Competing interests

No competing interests exist.

## Data sharing

All transcriptional datasets used in this manuscript were derived from public repositories. Their source is detailed in table S1 and software used to analyse these data is described in the methods.

## Funding

This work was supported by grant funding from the Wellcome Trust to MN (207511/Z/17/Z) and GP (WT101766/Z/13/Z), by Clinical Lectureship and Clinical Fellowship posts funded by the National Institute for Health Research (NIHR) to GP and LCKB respectively, and by NIH grants 5R01GM109018 and 5U54CA209997 to SDS. CEM would like to thank the Scientific Computing Unit (SCU), XSEDE Supercomputing Resources, the Starr Cancer Consortium (I13-0052), and funding from the WorldQuant Foundation, The Pershing Square Sohn Cancer Research Alliance, NASA (NNX14AH50G, NNX17AB26G), the National Institutes of Health (R21AI129851, R01MH117406, R01AI151059).

## Supplemental figure legends

**Figure S1.**
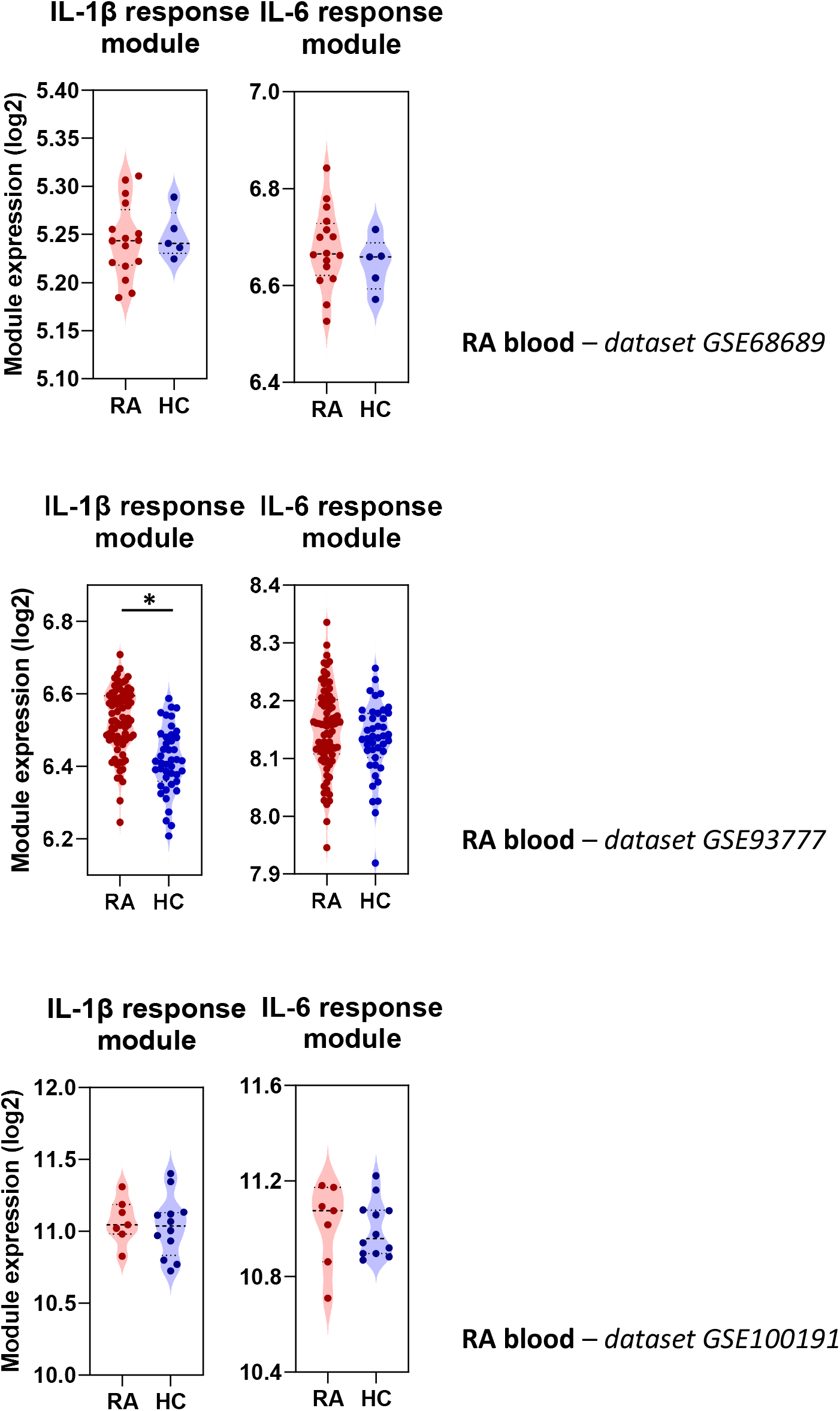
Cytokine response module expression in the blood of rheumatoid arthritis (RA) patients. Geometric mean expression of IL-1β and IL-6 cytokine response modules in the transcriptome of blood samples from RA patients compared to healthy controls. * = p < 0.05 by Mann-Whitney test. Transcriptomic datasets assessed are designated adjacent to each figure panel.

**Figure S2.**
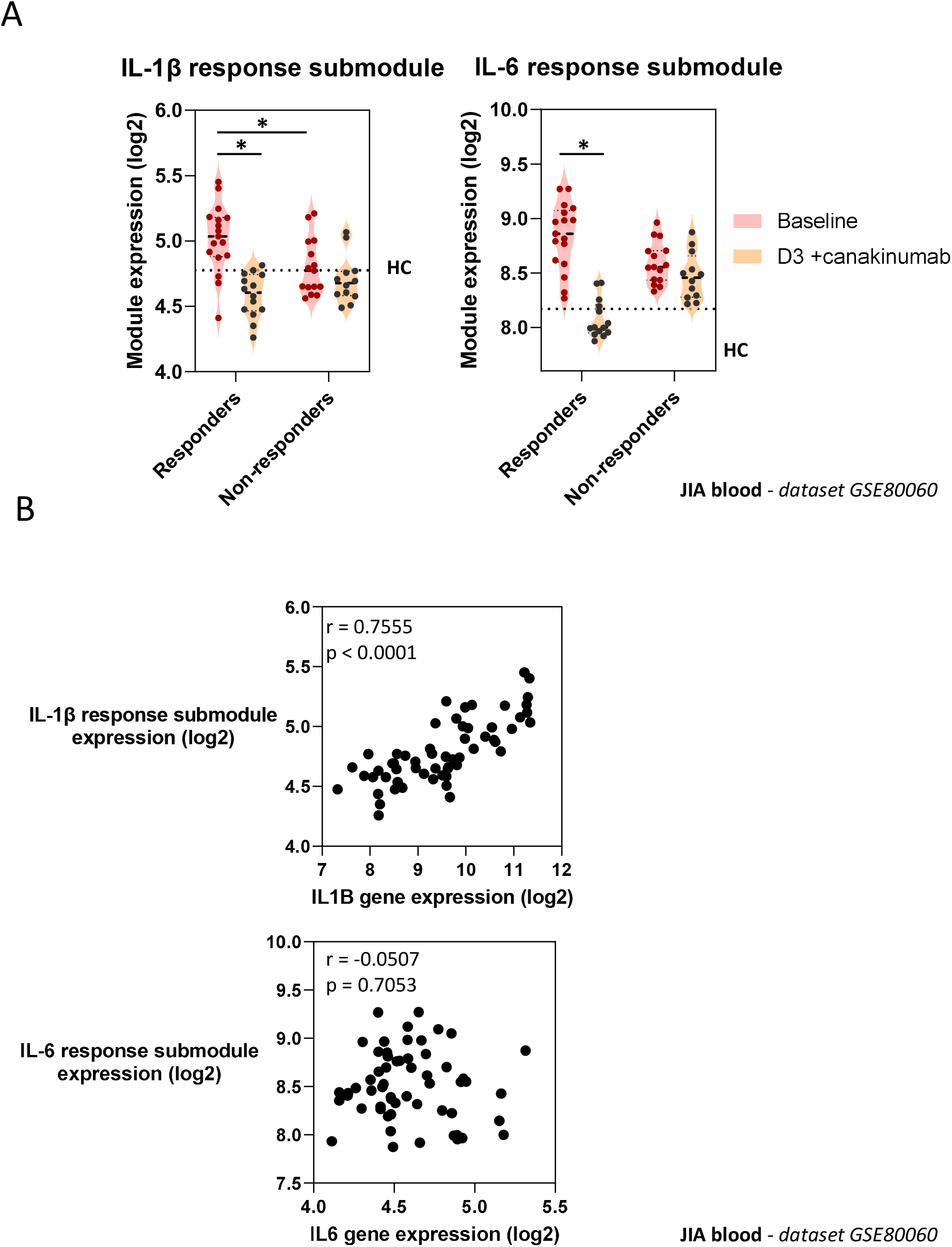
Effect of canakinumab on expression of cytokine genes and response submodules. A) Geometric mean expression of IL-1 and IL-6 cytokine response submodules in JIA patients before and 3 days after administration of canakinumab. Patients were subdivided into good responders (90-100% improvement) and non-responders (0-30% improvement). Dotted lines indicate median module or gene expression in healthy controls (HC) population in same dataset. * = p < 0.05 by Mann-Whitney test. B) Relationship between expression of cytokine response modules and cytokine genes. Statistical assessment of correlation made by Spearman Rank correlation. r = correlation coefficient. Transcriptomic dataset designated adjacent to figure panels.

**Figure S3.**
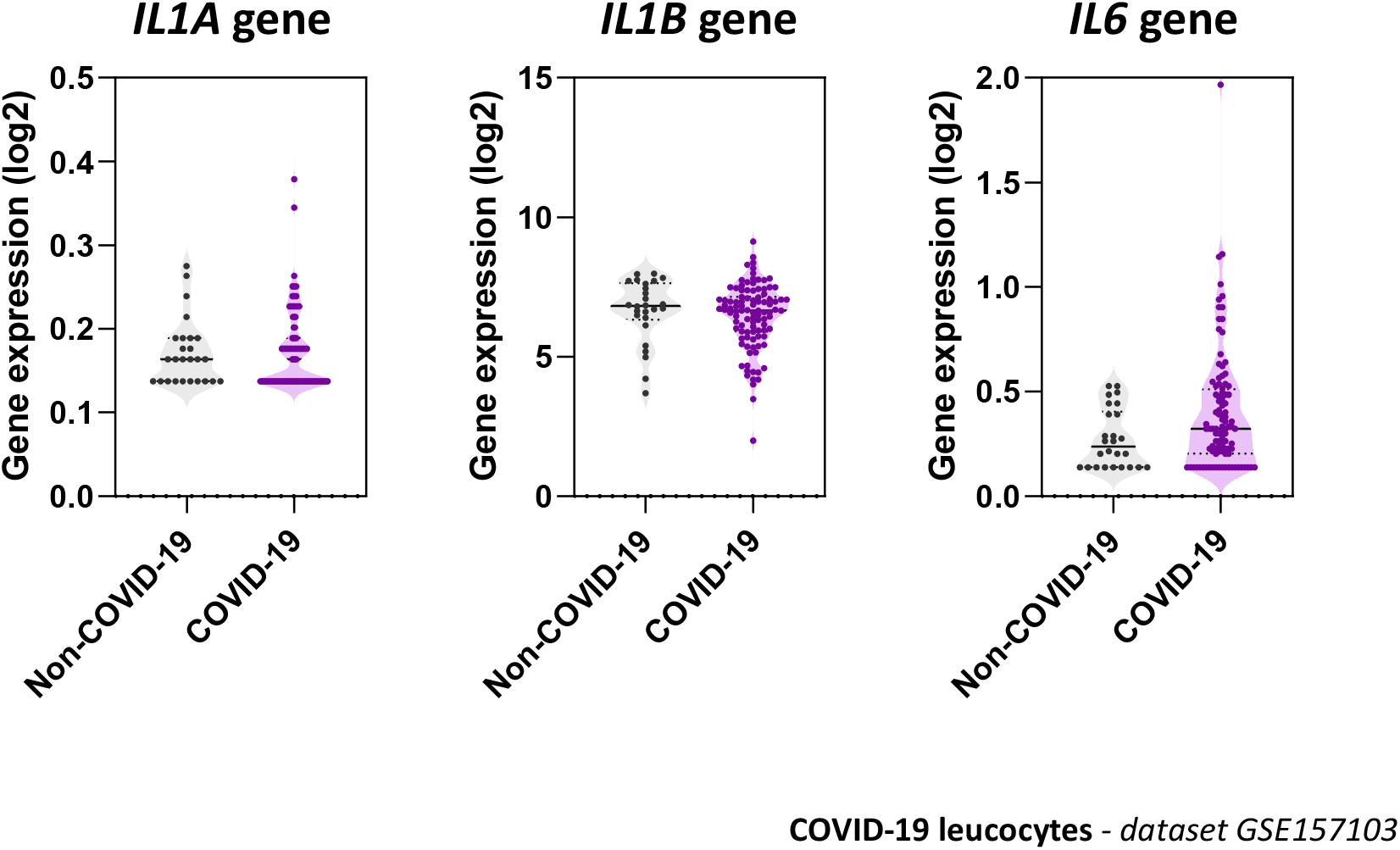
Cytokine gene expression in leucocytes of admitted patients with and without COVID-19. Expression of *IL1A*, *IL1B* and *IL6* genes in transcriptomic profiles of blood leucocytes collected from 101 COVID-19 and 24 non-COVID-19 patients. In this study, samples were collected from patients at a median of 3.37 days from admission to hospital (Overmyer et al., 2020). All comparisons were not significant by Mann-Whitney test. Transcriptomic dataset assessed are designated adjacent to each figure panel.

## Supplemental tables

**Table S1.**
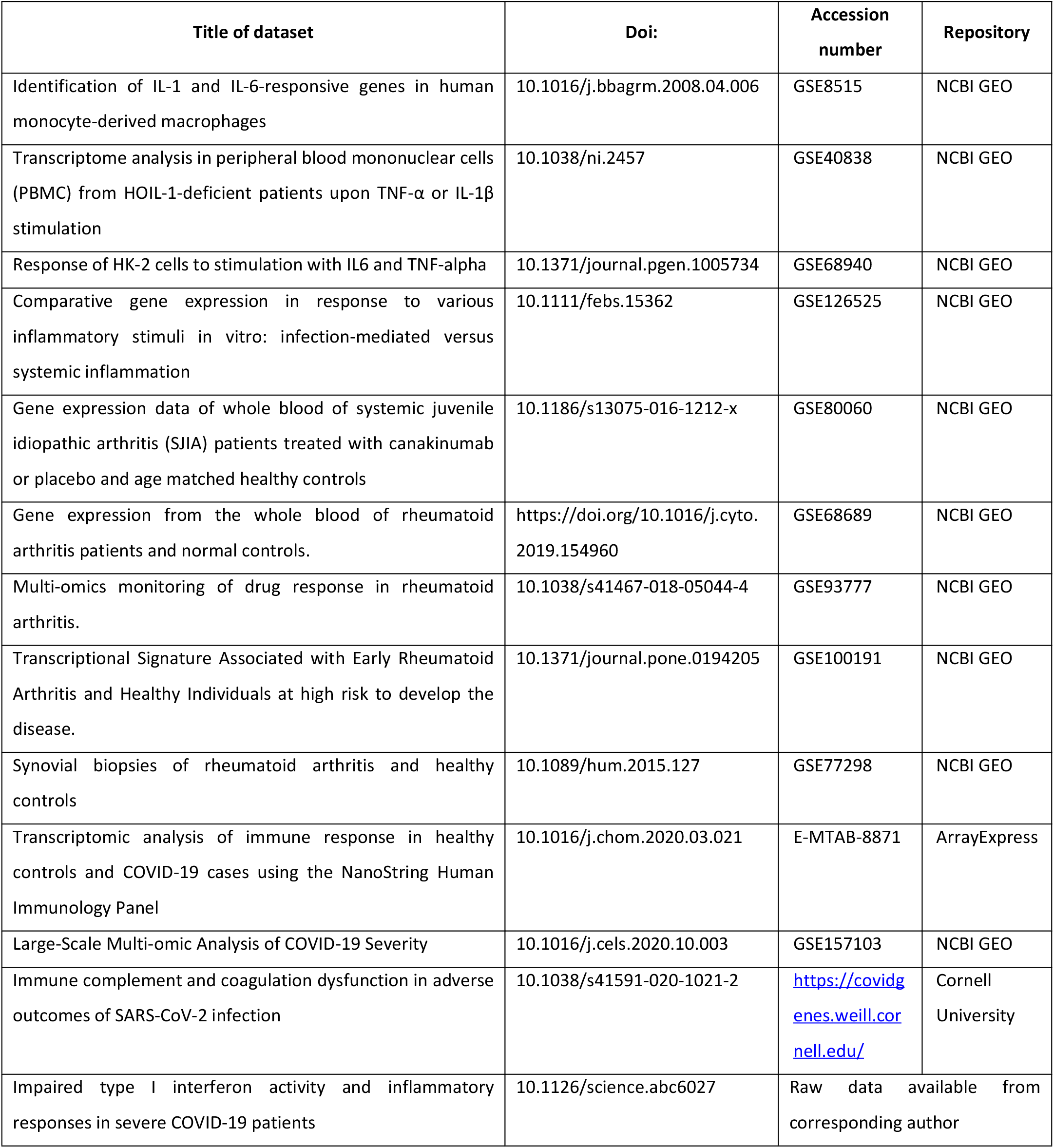
Transcriptional datasets used in this manuscript

**Table S2.** Cytokine response transcriptional modules and constituent genes

| Module name | Number of genes | Gene names |
| --- | --- | --- |
| **IL-1 response** | 57 | *ADORA2A, C15ORF48, C20ORF127, C2CD4B, C7ORF63, CCL20, CCL8, CHMP1B, CSF2, CSF3, CXCL1, CXCL2, CXCL5, CXCL6, DNAJB9, EGLN1, FGF2, FOXO3, GLIS3, GNA15, HAS3, HIATL1, IER3, IL11, IL6, ITPRIP, JARID2, KCNG1, LOC100134000, LRIG1, MAP3K8, MFSD2, MGC87042, MSL3, MT1G, MT1X, MTE, MTHFD2L, NAB1, NAMPT, NFKBIZ, NR4A2, OSGIN2, PFKFB3, PIM2, RCAN1, RNF145, SERPINB4, SGK1, SLC43A3, STEAP1, STEAP2, TFAP2C, TGIF1, TWIST2, ZC3H12A, ZC3H12C* |
| **IL-1 response submodule** | 7 | *CCL20, CCL8, CSF2, CXCL1, CXCL2, IL6, NFKBIZ* |
| **IL-6 response** | 41 | *AAMP, AKIP1, ANKRD10, ARPC2, CCR1, CD14, CSDE1, CTDSP2, CTNNA1, CXADR, DOK1, GADD45B, HAMP, IDH3B, IFI16, IL16, KAT5, LDLRAP1, MAP4, MR1, MSRB2, NCF4, NDUFB8, NPC1, PGD, PI4K2A, PPARD, PSMD4, RAP1GAP, RHOC, RIN2, RNASE1, RREB1, SASH1, SDS, SP110, STIP1, TSC22D3, UBE2M, UFL1, YBX3* |
| **IL-6 response submodule** | 7 | *CCR1, CD14, HAMP, IFI16, IL16, MR1, NCF4* |

## Notes

### Competing Interest Statement

The authors have declared no competing interest.

## References

Bell, L.C.K., Pollara, G., Pascoe, M., Tomlinson, G.S., Lehloenya, R.J., Roe, J., Meldau, R., Miller, R.F., Ramsay, A., Chain, B.M., et al. (2016). *In vivo* Molecular Dissection of the Effects of HIV-1 in Active Tuberculosis. PLoS Pathog. 12, e1005469.

De Benedetti, F., Brunner, H.I., Ruperto, N., Kenwright, A., Wright, S., Calvo, I., Cuttica, R., Ravelli, A., Schneider, R., Woo, P., et al. (2012). Randomized trial of tocilizumab in systemic juvenile idiopathic arthritis. N. Engl. J. Med. 367, 2385–2395.

Boisson, B., Laplantine, E., Prando, C., Giliani, S., Israelsson, E., Xu, Z., Abhyankar, A., Israël, L., Trevejo-Nunez, G., Bogunovic, D., et al. (2012). Immunodeficiency, autoinflammation and amylopectinosis in humans with inherited HOIL-1 and LUBAC deficiency. Nat. Immunol. 13, 1178–1186.

Brachat, A.H., Grom, A.A., Wulffraat, N., Brunner, H.I., Quartier, P., Brik, R., McCann, L., Ozdogan, H., Rutkowska-Sak, L., Schneider, R., et al. (2017). Early changes in gene expression and inflammatory proteins in systemic juvenile idiopathic arthritis patients on canakinumab therapy. Arthritis Res. Ther. 19, 13.

Broeren, M.G.A., de Vries, M., Bennink, M.B., Arntz, O.J., Blom, A.B., Koenders, M.I., van Lent, P.L.E.M., van der Kraan, P.M., van den Berg, W.B., and van de Loo, F.A.J. (2016). Disease-Regulated Gene Therapy with Anti-Inflammatory Interleukin-10 Under the Control of the CXCL10 Promoter for the Treatment of Rheumatoid Arthritis. Hum. Gene Ther. 27, 244–254.

Bullard, J., Dust, K., Funk, D., Strong, J.E., Alexander, D., Garnett, L., Boodman, C., Bello, A., Hedley, A., Schiffman, Z., et al. (2020). Predicting infectious SARS-CoV-2 from diagnostic samples. Clin. Infect. Dis.

Byng-Maddick, R., Turner, C.T., Pollara, G., Ellis, M., Guppy, N.J., Bell, L.C.K., Ehrenstein, M.R., and Noursadeghi, M. (2017). Tumor Necrosis Factor (TNF) Bioactivity at the Site of an Acute Cell-Mediated Immune Response Is Preserved in Rheumatoid Arthritis Patients Responding to Anti-TNF Therapy. Front. Immunol. 8, 932.

Das, A.S., Basu, A., Kumar, R., Borah, P.K., Bakshi, S., Sharma, M., Duary, R.K., Ray, P.S., and Mukhopadhyay, R. (2020). Post-transcriptional regulation of C-C motif chemokine ligand 2 expression by ribosomal protein L22 during LPS-mediated inflammation. FEBS J.

Dheda, K., Lenders, L., Srivastava, S., Magombedze, G., Wainwright, H., Raj, P., Bush, S.J., Pollara, G., Steyn, R., Davids, M., et al. (2019). Spatial Network Mapping of Pulmonary Multidrug-Resistant Tuberculosis Cavities Using RNA Sequencing. Am. J. Respir. Crit. Care Med.

Fleischmann, R. (2017). Interleukin-6 inhibition for rheumatoid arthritis. The Lancet 389, 1168–1170.

Gupta, R.K., Harrison, E.M., Ho, A., Docherty, A.B., Knight, S.R., van Smeden, M., Abubakar, I., Lipman, M., Quartagno, M., Pius, R.B., et al. (2020). Development and validation of the 4C Deterioration model for adults hospitalised with COVID-19. medRxiv.

Hadjadj, J., Yatim, N., Barnabei, L., Corneau, A., Boussier, J., Smith, N., Péré, H., Charbit, B., Bondet, V., Chenevier-Gobeaux, C., et al. (2020). Impaired type I interferon activity and inflammatory responses in severe COVID-19 patients. Science (80-.). 369, 718–724.

Hedrick, T.L., Murray, B.P., Hagan, R.S., and Mock, J.R. (2020). COVID-19: Clean up on IL-6. Am. J. Respir. Cell Mol. Biol.

Hermine, O., Mariette, X., Tharaux, P.-L., Resche-Rigon, M., Porcher, R., Ravaud, P., and CORIMUNO-19 Collaborative Group (2020). Effect of Tocilizumab vs Usual Care in Adults Hospitalized With COVID-19 and Moderate or Severe Pneumonia: A Randomized Clinical Trial. JAMA Intern. Med.

Huang, C., Wang, Y., Li, X., Ren, L., Zhao, J., Hu, Y., Zhang, L., Fan, G., Xu, J., Gu, X., et al. (2020). Clinical features of patients infected with 2019 novel coronavirus in Wuhan, China. The Lancet 395, 497–506.

Iwasaki, H., Takeuchi, O., Teraguchi, S., Matsushita, K., Uehata, T., Kuniyoshi, K., Satoh, T., Saitoh, T., Matsushita, M., Standley, D.M., et al. (2011). The IκB kinase complex regulates the stability of cytokine-encoding mRNA induced by TLR-IL-1R by controlling degradation of regnase-1. Nat. Immunol. 12, 1167–1175.

Jura, J., Wegrzyn, P., Korostyński, M., Guzik, K., Oczko-Wojciechowska, M., Jarzab, M., Kowalska, M., Piechota, M., Przewłocki, R., and Koj, A. (2008). Identification of interleukin-1 and interleukin-6-responsive genes in human monocyte-derived macrophages using microarrays. Biochim. Biophys. Acta 1779, 383–389.

Lee, E.J., Lilja, S., Li, X., Schäfer, S., Zhang, H., and Benson, M. (2020). Bulk and single cell transcriptomic data indicate that a dichotomy between inflammatory pathways in peripheral blood and arthritic joints complicates biomarker discovery. Cytokine 127, 154960.

Leisman, D.E., Ronner, L., Pinotti, R., Taylor, M.D., Sinha, P., Calfee, C.S., Hirayama, A.V., Mastroiani, F., Turtle, C.J., Harhay, M.O., et al. (2020). Cytokine elevation in severe and critical COVID-19: a rapid systematic review, meta-analysis, and comparison with other inflammatory syndromes. Lancet Respir. Med.

Liao, M., Liu, Y., Yuan, J., Wen, Y., Xu, G., Zhao, J., Cheng, L., Li, J., Wang, X., Wang, F., et al. (2020). Single-cell landscape of bronchoalveolar immune cells in patients with COVID-19. Nat. Med. 26, 842–844.

Macías-Segura, N., Castañeda-Delgado, J.E., Bastian, Y., Santiago-Algarra, D., Castillo-Ortiz, J.D., Alemán-Navarro, A.L., Jaime-Sánchez, E., Gomez-Moreno, M., Saucedo-Toral, C.A., Lara-Ramírez, E.E., et al. (2018). Transcriptional signature associated with early rheumatoid arthritis and healthy individuals at high risk to develop the disease. PLoS One 13, e0194205.

Maes, B., Bosteels, C., De Leeuw, E., Declercq, J., Van Damme, K., Delporte, A., Demeyere, B., Vermeersch, S., Vuylsteke, M., Willaert, J., et al. (2020). Treatment of severely ill COVID-19 patients with anti-interleukin drugs (COV-AID): A structured summary of a study protocol for a randomised controlled trial. Trials 21, 468.

McGonagle, D., O’Donnell, J.S., Sharif, K., Emery, P., and Bridgewood, C. (2020). Immune mechanisms of pulmonary intravascular coagulopathy in COVID-19 pneumonia. The Lancet Rheumatology 2, e437–e445.

Mehta, P., McAuley, D.F., Brown, M., Sanchez, E., Tattersall, R.S., Manson, J.J., and HLH Across Speciality Collaboration, UK (2020). COVID-19: consider cytokine storm syndromes and immunosuppression. The Lancet 395, 1033–1034.

Nikfar, S., Saiyarsarai, P., Tigabu, B.M., and Abdollahi, M. (2018). Efficacy and safety of interleukin-1 antagonists in rheumatoid arthritis: a systematic review and meta-analysis. Rheumatol. Int. 38, 1363–1383.

Nishimoto, N., Terao, K., Mima, T., Nakahara, H., Takagi, N., and Kakehi, T. (2008). Mechanisms and pathologic significances in increase in serum interleukin-6 (IL-6) and soluble IL-6 receptor after administration of an anti-IL-6 receptor antibody, tocilizumab, in patients with rheumatoid arthritis and Castleman disease. Blood 112, 3959–3964.

O’Brown, Z.K., Van Nostrand, E.L., Higgins, J.P., and Kim, S.K. (2015). The Inflammatory Transcription Factors NFκB, STAT1 and STAT3 Drive Age-Associated Transcriptional Changes in the Human Kidney. PLoS Genet. 11, e1005734.

Ong, E.Z., Chan, Y.F.Z., Leong, W.Y., Lee, N.M.Y., Kalimuddin, S., Haja Mohideen, S.M., Chan, K.S., Tan, A.T., Bertoletti, A., Ooi, E.E., et al. (2020). A Dynamic Immune Response Shapes COVID-19 Progression. Cell Host Microbe 27, 879–882.e2.

Overmyer, K.A., Shishkova, E., Miller, I.J., Balnis, J., Bernstein, M.N., Peters-Clarke, T.M., Meyer, J.G., Quan, Q., Muehlbauer, L.K., Trujillo, E.A., et al. (2020). Large-Scale Multi-omic Analysis of COVID-19 Severity. Cell Syst.

Pollara, G., Murray, M.J., Heather, J.M., Byng-Maddick, R., Guppy, N., Ellis, M., Turner, C.T., Chain, B.M., and Noursadeghi, M. (2017). Validation of immune cell modules in multicellular transcriptomic data. PLoS One 12, e0169271.

Pollara, G., Turner, C.T., Tomlinson, G.S., Bell, L.C., Khan, A., Peralta, L.F., Folino, A., Akarca, A., Venturini, C., Baker, T., et al. (2019). Exaggerated *in vivo* IL-17 responses discriminate recall responses in active TB. BioRxiv.

Qin, C., Zhou, L., Hu, Z., Zhang, S., Yang, S., Tao, Y., Xie, C., Ma, K., Shang, K., Wang, W., et al. (2020). Dysregulation of Immune Response in Patients With Coronavirus 2019 (COVID-19) in Wuhan, China. Clin. Infect. Dis. 71, 762–768.

Ramlall, V., Thangaraj, P.M., Meydan, C., Foox, J., Butler, D., Kim, J., May, B., De Freitas, J.K., Glicksberg, B.S., Mason, C.E., et al. (2020). Immune complement and coagulation dysfunction in adverse outcomes of SARS-CoV-2 infection. Nat. Med. 26, 1609–1615.

Ravindra, N.G., Alfajaro, M.M., Gasque, V., Wei, J., Filler, R.B., Huston, N.C., Wan, H., Szigeti-Buck, K., Wang, B., Montgomery, R.R., et al. (2020). Single-cell longitudinal analysis of SARS-CoV-2 infection in human bronchial epithelial cells. BioRxiv.

Ruperto, N., Brunner, H.I., Quartier, P., Constantin, T., Wulffraat, N., Horneff, G., Brik, R., McCann, L., Kasapcopur, O., Rutkowska-Sak, L., et al. (2012). Two randomized trials of canakinumab in systemic juvenile idiopathic arthritis. N. Engl. J. Med. 367, 2396–2406.

Salvarani, C., Dolci, G., Massari, M., Merlo, D.F., Cavuto, S., Savoldi, L., Bruzzi, P., Boni, F., Braglia, L., Turrà, C., et al. (2020). Effect of Tocilizumab vs Standard Care on Clinical Worsening in Patients Hospitalized With COVID-19 Pneumonia: A Randomized Clinical Trial. JAMA Intern. Med.

Seko, Y., Cole, S., Kasprzak, W., Shapiro, B.A., and Ragheb, J.A. (2006). The role of cytokine mRNA stability in the pathogenesis of autoimmune disease. Autoimmun Rev. 5, 299–305.

Stone, J.H., Frigault, M.J., Serling-Boyd, N.J., Fernandes, A.D., Harvey, L., Foulkes, A.S., Horick, N.K., Healy, B.C., Shah, R., Bensaci, A.M., et al. (2020). Efficacy of Tocilizumab in Patients Hospitalized with Covid-19. N. Engl. J. Med.

Subramanian, A., Tamayo, P., Mootha, V.K., Mukherjee, S., Ebert, B.L., Gillette, M.A., Paulovich, A., Pomeroy, S.L., Golub, T.R., Lander, E.S., et al. (2005). Gene set enrichment analysis: a knowledge-based approach for interpreting genome-wide expression profiles. Proc. Natl. Acad. Sci. USA 102, 15545–15550.

Tasaki, S., Suzuki, K., Kassai, Y., Takeshita, M., Murota, A., Kondo, Y., Ando, T., Nakayama, Y., Okuzono, Y., Takiguchi, M., et al. (2018). Multi-omics monitoring of drug response in rheumatoid arthritis in pursuit of molecular remission. Nat Commun 9, 2755.

Thwaites, R., Sanchez Sevilla Uruchurtu, A., Siggins, M., Liew, F., Russell, C.D., Moore, S., Carter, E., Abrams, S., Short, C.-E., Thaventhiran, T., et al. (2020). Elevated antiviral, myeloid and endothelial inflammatory markers in severe COVID-19. medRxiv.

Zhang, X., Tan, Y., Ling, Y., Lu, G., Liu, F., Yi, Z., Jia, X., Wu, M., Shi, B., Xu, S., et al. (2020). Viral and host factors related to the clinical outcome of COVID-19. Nature 583, 437–440.

Zhou, F., Yu, T., Du, R., Fan, G., Liu, Y., Liu, Z., Xiang, J., Wang, Y., Song, B., Gu, X., et al. (2020a). Clinical course and risk factors for mortality of adult inpatients with COVID-19 in Wuhan, China: a retrospective cohort study. The Lancet 395, 1054–1062.

Zhou, Z., Ren, L., Zhang, L., Zhong, J., Xiao, Y., Jia, Z., Guo, L., Yang, J., Wang, C., Jiang, S., et al. (2020b). Overly Exuberant Innate Immune Response to SARS-CoV-2 Infection. SSRN Journal.

